# Dynamics of auditory word form encoding in human speech cortex

**DOI:** 10.1101/2025.05.05.651964

**Authors:** Yizhen Zhang, Matthew K. Leonard, Laura Gwilliams, Ilina Bhaya-Grossman, Edward F. Chang

**Affiliations:** Department of Neurological Surgery, University of California, San Francisco, San Francisco, CA, USA; Weill Institute for Neuroscience, University of California, San Francisco, San Francisco, CA, USA; University of California, Berkeley - University of California, San Francisco Graduate Program in Bioengineering, Berkeley, CA, USA

## Abstract

When we hear continuous speech, we perceive it as a series of discrete words, despite the lack of clear boundaries in the acoustic signal. The superior temporal gyrus (STG) encodes phonetic elements like consonants and vowels, but how it extracts whole words as perceptual units remains unclear. Using high-density cortical recordings, we investigated how the brain represents auditory word forms—integrating acoustic-phonetic, prosodic, and lexical features—while participants listened to spoken narratives. Our results show that STG neural populations exhibit a distinctive reset in activity at word boundaries, marked by a brief, sharp drop in cortical activity. Between these resets, the STG consistently encodes distinct acoustic-phonetic, prosodic, and lexical information, supporting the integration of phonological features into coherent word forms. Notably, this process tracks the relative elapsed time within each word, independent of its absolute duration, providing a flexible temporal scaffolding for encoding variable word lengths. We observed similar word form dynamics in the deeper layers of a self-supervised artificial speech network, suggesting a potential convergence with computational models. Additionally, in a bistable word perception task, STG responses were aligned with participants’ perceived word boundaries on a trial-by-trial basis, further emphasizing the role of dynamic encoding in word recognition. Together, these findings support a new dynamical model of auditory word forms, highlighting their importance as perceptual units for accessing linguistic meaning.

## Introduction

In 1874, the neurologist Karl Wernicke observed a patient who lost the ability to comprehend speech after injury to the left superior temporal gyrus (STG), now called Wernicke’s area. He proposed that this region processes the “auditory sensory image of words”, referring to the holistic perception of word units (Wernicke, 1874). While recent research has decoded how the brain processes speech sounds - such as consonants, vowels, and prosodic cues like pitch and rhythm (Leonard et al., 2024; Mesgarani et al., 2014; Oganian et al., 2023; Oganian & Chang, 2019; Tang et al., 2017) - how these sounds are integrated into the perceptual units we hear, words, remains unclear and debated.

The auditory word form is a key perceptual unit that combines acoustic-phonetic features (consonants and vowels), prosodic elements (pitch, intensity, rhythm), and lexical information (word frequency and duration) into a cohesive whole recognized by the brain as a specific word (Binder, 2015; Norris et al., 1997). One prevailing theory suggests that sub-lexical phonological content encoded in the STG is transformed into word-level representations by higher-order brain regions like the anterior temporal cortex, posterior middle temporal gyrus (MTG), or inferior frontal gyrus (IFG) (Damera et al., 2023; DeWitt & Rauschecker, 2012; Jasmin et al., 2019; Wilson et al., 2018). However, evidence for the precise neural representation of word units has been inconsistent, partly due to limitations in spatial and temporal resolution of imaging techniques.

Furthermore, given the rapid and variable nature of spoken language, a largely serial feedforward view that does not take into account dynamics may ultimately prove insufficient to understand how the brain maps sequences of sounds onto word representations (Bhaya-Grossman & Chang, 2022). An alternative approach emphasizes how neural activity integrates information over time rather than across distinct brain regions (W. D. Marslen-Wilson, 1987; W. Marslen-Wilson & Tyler, 1980; Yi et al., 2019). By focusing on neural activity aligned to the events around which words are organized - word boundaries - in continuous speech, we can directly examine how the brain represents auditory word forms.

To explore this, we used high-density electrocorticography (ECoG) to record neural activity with millimeter and millisecond precision as participants listened to continuous, natural speech. This allowed us to track single-trial neural activity during real-time language comprehension. We identified STG neural populations that showed distinct neurophysiological response patterns aligned to word boundaries, as well as encoding of acoustic-phonetic, prosodic, and lexical features between these boundaries.

We applied state-space modeling of the STG population trajectory, and found a temporal scaffolding mechanism that tracks word duration and relative timing within words. Similar dynamics were observed in the deeper layers of a self-supervised deep learning model of speech, providing convergence with representations learned by an artificial computational model. Finally, in a bistable word perception task, neural responses in the STG aligned with participants’ subjective perception of word forms, demonstrating the role of these populations in shaping perceptual outcomes.

Together, these findings provide compelling evidence for a dynamic, integrative neural mechanism in the STG that underlies the representation of auditory word forms in continuous speech.

## Results

### A neurophysiological marker of word boundaries in continuous speech

The acoustic cues that can indicate the location of word boundaries in continuous speech - e.g., pauses, specific phonemes, and prosodic cues like stress and duration - are highly variable and unreliable in natural speech (**Fig. 1A**) (Lehiste, 1972). Thus, it is currently unknown whether neural populations show specific activity aligned to spoken words. A key question is whether the human brain distinguishes syllables, which are marked by rapid changes in the speech amplitude envelope (intensity (Oganian & Chang, 2019)), and words, which are a subset of all syllable boundaries in spoken English (**Fig. 1B**).

**Fig. 1:**
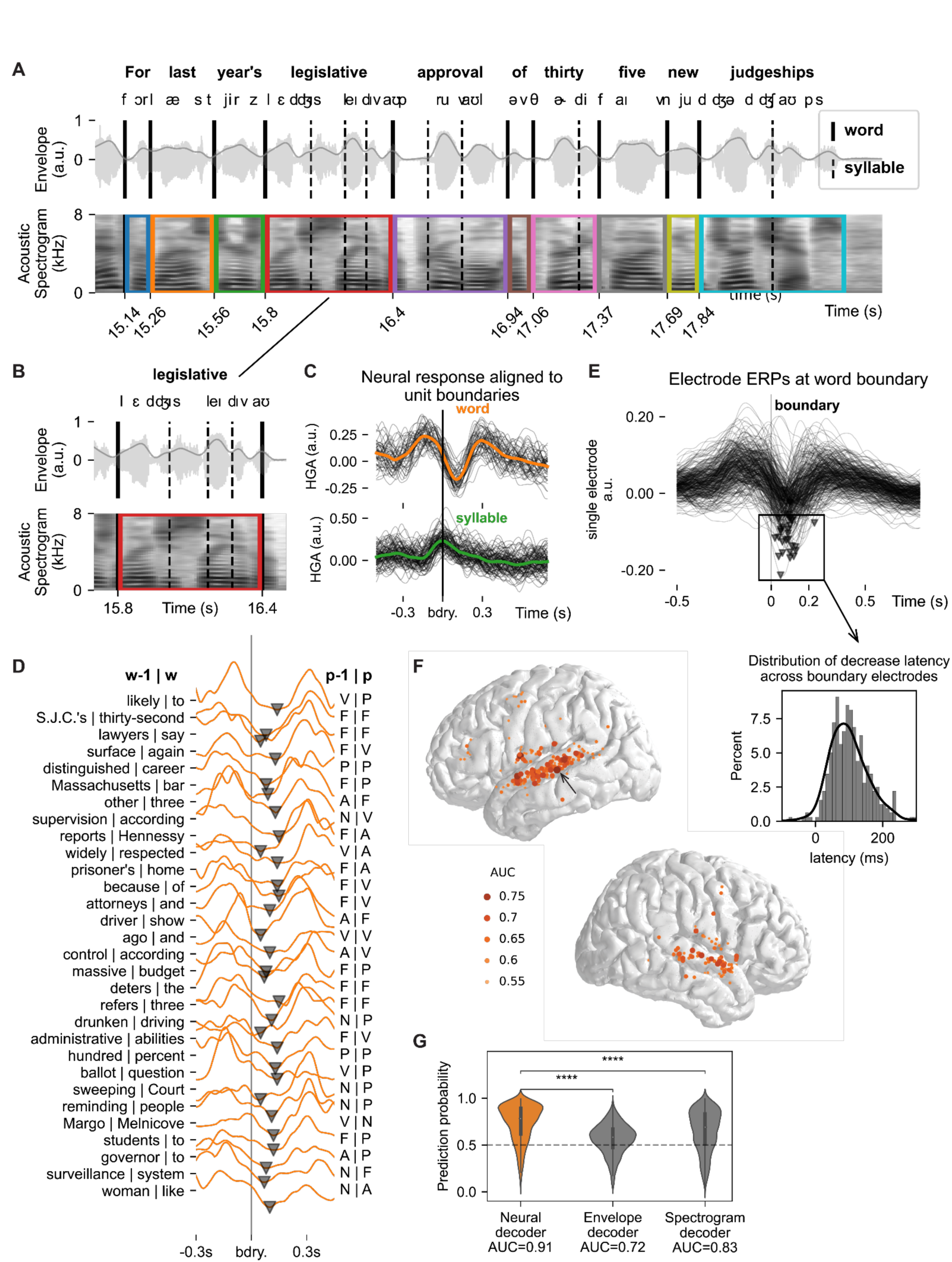
Neural populations mark word boundaries in natural continuous speech. A. An example spoken phrase illustrates the challenge in identifying word boundaries in continuous speech. From top to bottom: sentence, phoneme label, acoustic waveform, envelope, and spectrogram. Word boundaries and syllable boundaries are marked by solid and dashed vertical lines, respectively. Colored segments over the spectrogram show speech corresponding to auditory word forms, which are not consistently marked by acoustic cues. B. Example of the ambiguous acoustic cues to word and syllable boundaries in the multi-syllabic word “legislative”. Both syllables and words are marked by similar cues in the waveform, envelope, and spectrogram. C. Neural activity aligned to word (top) and syllable (bottom) boundaries has distinct temporal patterns for an example electrode in STG (see arrow in F), suggesting that neural populations have unique word-aligned activity, despite ambiguous acoustic cues. Word boundaries are uniquely marked by a transient decrease in high-gamma amplitude (HGA) immediately after the boundary. D. The transient decrease in HGA is observed on example single trials aligned to word boundaries for the example electrode in C. Example trials are annotated with the preceding (w-1) and current (w) words, and the preceding (p-1) and current (p) acoustic-phonetic features (V: vowel, P: plosive, F: fricative, N: nasal, A: approximant). The same neural activity pattern is observed regardless of the specific speech content. E. Top: Word boundary-aligned neural activity for all electrodes with significant word boundary decoding (see Methods). Bottom: Distribution of boundary decrease latency (mean = 99.1ms, standard deviation = 65.5ms) across all electrodes with significant word boundary decoding. F. Electrodes with significant word boundary decoding are localized primarily to bilateral STG. AUC indicates area under the curve for the word versus syllable boundary classifier. G. Neural activity in STG is a better predictor of word boundaries than acoustics. Violin plots show distributions of prediction probabilities from decoders trained on all boundary electrodes (orange, left), amplitude envelope (gray, middle), and spectrogram (gray, right).

Participants (n=16) passively listened to 10 short (181s±29s) spoken narratives from radio news stories spoken by news announcers (Boston University Radio News Corpus) (Ostendorf et al., 1995). These audio recordings were annotated for within-word syllables and word boundaries (see Methods), along with phonemes and prosodic markers.

First, we asked whether neural activity changes at word boundaries even though such boundaries are ambiguously cued by the acoustic input. We identified electrodes where a sharp, transient decrease in the power of the high-gamma amplitude (HGA; 70-150Hz) occurred consistently ∼100ms after word boundaries (**Fig. 1C**, top). This decrease was unique to word boundaries, since within-word syllables exhibited an increased response, as observed in prior work (Oganian & Chang, 2019) (**Fig. 1C**, bottom). The word boundary-specific decrease was apparent on single trials where the characteristics of the two words at the boundary (word durations, stress patterns, last/first phonemes, etc) were highly variable (**Fig. 1D**). These results demonstrate that the human brain tracks the boundaries between words in continuous natural speech.

Across participants, we identified 321 electrodes that significantly differentiated word boundaries from syllable boundaries (logistic regression classifiers with L2 regularization; permutation test, one-sided p<0.001). Of these, 319 (99.4%) had a prominent decrease (peak prominence>0.05, peak width>50ms; see Methods) in HGA immediately after word boundaries (**Fig. 1E**; mean ± standard deviation of boundary peak latency: 99.1±65.5ms). These word boundary electrodes were located primarily in bilateral STG, spanning a majority of the gyrus from posterior to mid-anterior sub-regions (**Fig. 1F**). There were also some electrodes on the ventral pre-central gyrus (PreCG) and left inferior frontal gyrus (IFG), which are known to have auditory responses as part of a parallel speech pathway (Hullett et al., 2024).

To understand to what extent these neural populations rely on information beyond the (unreliable) acoustic cues to identify word boundaries in natural speech, we trained a logistic regression classifier using (1) the population of electrodes with significant word boundary decoding; (2) the acoustic speech envelope; and (3) the acoustic spectrogram of speech. We found that the decoder trained using neural data demonstrated significantly higher performance (**Fig. 1G**; neural decoder AUC=0.91; envelope decoder AUC=0.72; spectrogram decoder AUC=0.83; paired t-test across trials, p<0.0001). Together, these results demonstrate that neural populations in STG track word boundaries in continuous speech, above and beyond the acoustic cues that correlate with this crucial perceptual and linguistic unit.

### Acoustic-phonetic, prosodic, and lexical encoding aligned to word boundaries

Having demonstrated that neural populations in STG track the boundaries between critical perceptual units of speech - words - we next asked how various features of the words themselves are encoded in the neural activity between word boundaries. Based on prior studies (Chomsky and Halle 1991; Leonard et al. 2024; Mesgarani et al. 2014; Miller and Eimas 1995; Stevens 2002; McClelland and Elman 1986; Cole and Jakimik 1978; Marslen-Wilson and Welsh 1978), we selected a large set of features that we hypothesized could modulate neural activity and contribute to encoding of auditory word forms (**Fig. 2A**; see Methods for a detailed description of all features). To understand how these features are encoded around timepoints surrounding word boundaries, we applied a sliding-window partial correlation analysis (see Methods) to electrodes for which there were clear word-boundary evoked response patterns.

**Fig. 2:**
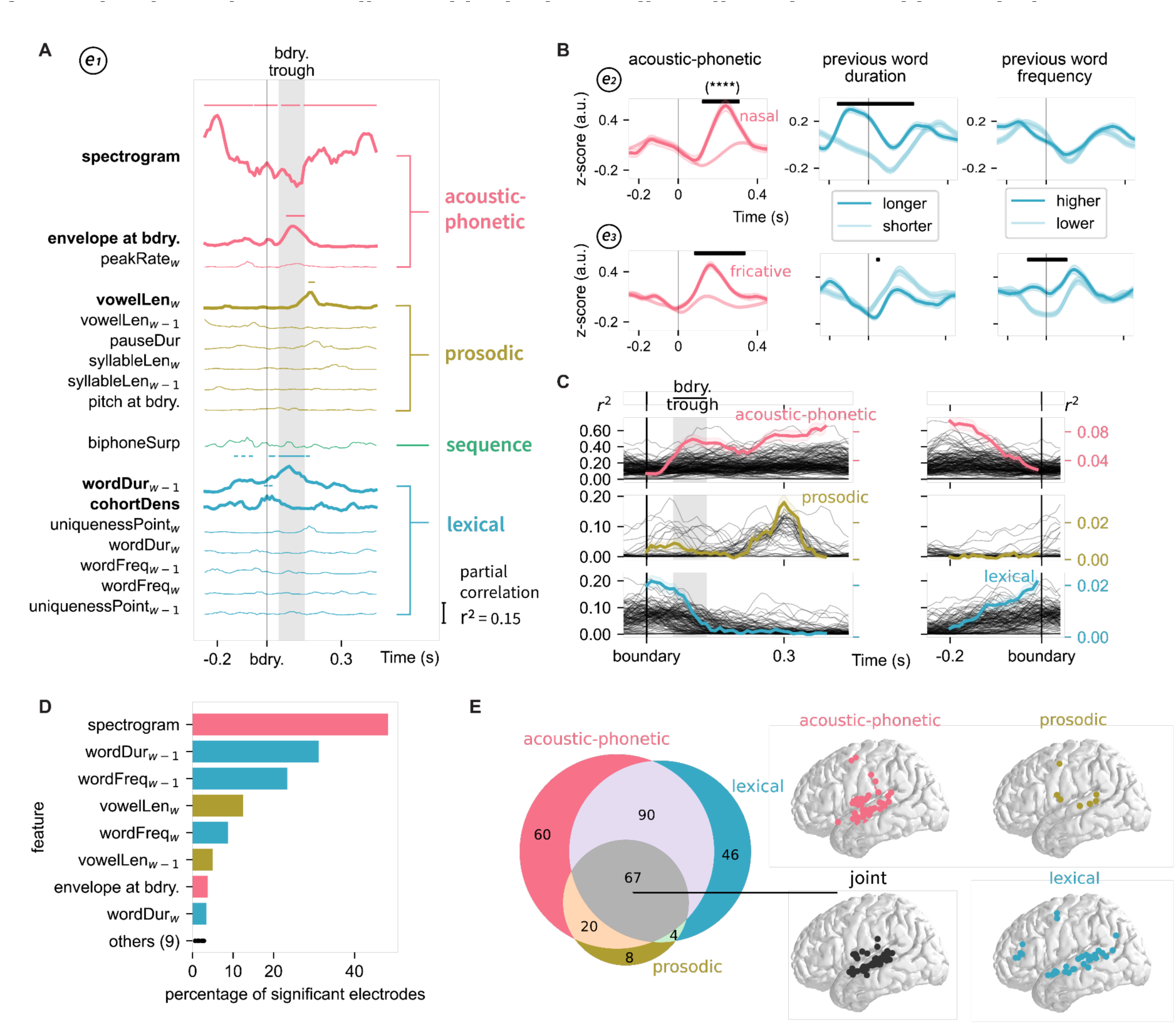
Neural activity in local STG populations between word boundaries dynamically encodes phonological features and characteristics of whole word forms. A. An example STG electrode with a clear word boundary-evoked activity pattern (top), is modulated by several distinct types of features at different latencies relative to the boundary (bottom; partial r^2^, p<0.001, Bonferroni correction; significant features are highlighted in bold text and with thick lines). B. Word boundary-evoked HGA differentiates specific phonological and word-level features. Example electrodes illustrate word-initial nasals (e2) and fricatives (e3; dark lines; left column) from all other acoustic-phonetic features (light lines). Other electrodes are also modulated by the duration of the previous word (middle column) and/or the frequency of the previous word (right column). Black bars indicate significant differences between bins of trials (one-way anova, p<0.0001). C. Neural activity between word boundaries is modulated by different classes of features at different times. Acoustic-phonetic encoding occurs throughout the word, except immediately around the boundary (top). Prosodic features show effects transiently ∼300ms after word onset, overlapping with acoustic-phonetic encoding (middle). Finally, lexical features are encoded primarily in the window surrounding the word boundary. Black lines indicate individual electrodes, colored lines indicate the mean across electrodes (right y-axis). Data are shown aligned to both the current word (left) and next word boundaries (right). D. Specific features reflected in the feature class averages in B-C, sorted by percentage of electrodes with significant effects. Colors indicate the feature class. E. Neural populations encoding acoustic-phonetic, prosodic, and lexical features around word boundaries overlap in mid-STG. A majority of electrodes encode more than one type of feature. The Venn diagram illustrates the number of significant electrodes.

As described above, neural activity time-locked to word boundaries is characterized by rapid, transient decreases in HGA (**Fig. 1C-E**). For an example electrode, we found that acoustic-phonetic, prosodic, and lexical features significantly modulate this activity, with different features showing unique encoding at different processing stages relative to word onset (**Fig. 2A**; partial r^2^, p<0.001, Bonferroni corrected). This example electrode also demonstrates that, while not all individual features are encoded within a single neural population, single electrode HGA can show significant modulation from multiple classes of speech and language features spanning the auditory and linguistic hierarchy.

To illustrate how this modulation manifests in HGA around word boundaries, we examined additional example electrodes. We found electrodes that showed significant differences for particular acoustic-phonetic features at word onset, for example nasals (e2) and fricatives (e3; **Fig. 2B**, left column; one-way anova, p<0.0001). These same electrodes could also show significant modulation from word-level features like the duration of the previous word (middle column) and/or the frequency of the previous word (right column). These examples suggest that individual neural populations in STG show activity aligned to word boundaries that is modulated by multiple features with distinct time courses.

Next, we asked how encoding of these features occurs over the duration of individual words across all electrodes with significant word boundary decoding. We examined the time course of partial r^2^ for each feature class, and found that across electrodes, there is a consistent sequence of encoding relative to the word boundary (**Fig. 2C**). Specifically, acoustic-phonetic features are encoded starting ∼100ms after the boundary (during the time of the transient decrease in the evoked HGA), and continuing until ∼100ms before the next word. Overlapping with acoustic-phonetic features, prosodic features show a transient encoding peak at ∼300ms after the word boundary. Finally, lexical features complement the timing of acoustic-phonetic encoding, showing significant partial r^2^ in the ∼200ms around the word boundary.

We identified a specific subset of acoustic-phonetic, prosodic, and lexical features that significantly explained neural activity aligned to words. These included spectrotemporal content (McClelland & Elman, 1986), markers of stress such as vowel length (Cutler 2008), and characteristics of the whole word form such as word frequency and word duration (Connine et al., 1993) (**Fig. 2D**). Together, these results demonstrate that neural populations throughout the STG are modulated by a highly diverse set of features that span the speech hierarchy, with timing that is specifically organized around words in continuous speech.

Finally, we asked whether distinct neural populations or brain regions encode different types of information associated with word-forms, as is predicted by dominant models of speech processing (DeWitt & Rauschecker, 2012; Hickok & Poeppel, 2007). We identified whether each electrode had significant encoding for any features in each class (acoustic-phonetic, prosodic, and lexical), and characterized the distribution of electrode types. We found that 38.6% (n=114) of electrodes encoded only one type of feature, with the majority being tuned to acoustic-phonetic features (**Fig. 2E**, left). Of the remaining electrodes that encoded more than one feature (61.4%, n=181), 38.6% (n=114) significantly encoded two features, and 22.7% (n=67) significantly encoded all three. The electrodes encoding all three features were located primarily in the mid-STG, a region that also showed populations that uniquely encoded each feature. These results demonstrate that the neural encoding of word forms is strongly associated with a spatially overlapping representation of multiple types of phonological speech information at the sub-lexical and lexical levels, with dynamic encoding of these features across time.

### Temporal tracking of word dynamics in neural populations

Thus far, we have demonstrated that auditory word forms are encoded in STG neural populations as the integrated set of acoustic-phonetic, prosodic, and lexical features between word boundaries. It remains unclear how these neural populations track when each word begins and ends, and therefore how the brain implements consistent, relative temporal patterns of feature encoding (**Fig. 2**) regardless of individual word duration. This is particularly challenging given that words are highly variable in duration (30-1310 ms in the stimulus used here), requiring a mechanism to represent the auditory word form information within the temporal framework of words unfolding in continuous speech.

We hypothesized that population neural activity tracks how much of a word has been heard in a continuous fashion, in other words, the relative elapsed time since a word boundary. To test this, we examined the time-course of neural activity for groups of words binned by duration. We found that neural activity in boundary electrodes scaled with duration, both before and during the transient decrease in HGA that marks word boundaries (**Fig. 3A**). Across electrodes, word duration could be decoded significantly above chance from this activity (26.2% of individual electrodes; r = 0.119±0.047, p<0.001, Mann-Whitney-Wilcoxon test, 10-fold cross-validation, **Fig. 3A**, bottom). This result suggests that by the end of a word in continuous speech, neural populations encode the amount of time that has elapsed since the word began.

**Fig. 3:**
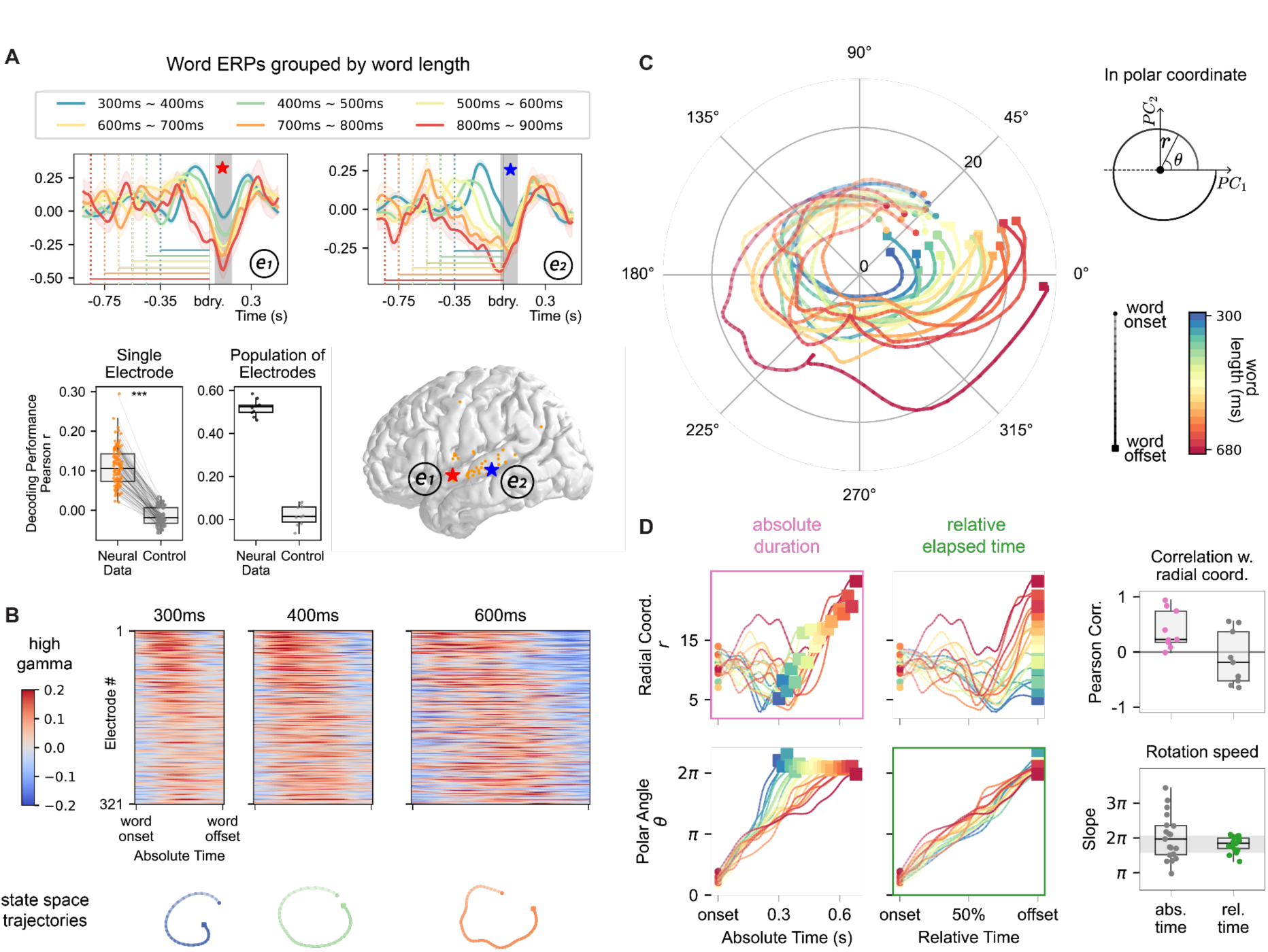
Neural population dynamics track word duration and relative elapsed time since word onset. A. Top: Word-locked HGA grouped by word duration (bin width = 100ms) in two example word boundary electrodes. HGA around word boundaries is correlated with the duration of the previous word. Bottom: Word duration can be decoded significantly above chance (control: permutation test, Mann-Whitney-Wilcoxon test, 10-fold cross-validation, ***:p<0.001) from the transient decrease in HGA (shaded gray in top panel) using single electrodes (left) or all (right) boundary electrodes. B. Top: HGA for all word boundary electrodes, averaged over words with 300ms, 400ms, or 600ms durations. Bottom: Projecting population-level neural activity for each binned word duration into the first two principal components (PCs) illustrates that neural trajectory dynamics reflect a complete cycle, regardless of word duration. C. All trajectories of population-level neural responses to words, binned by word duration (20ms bin size) in a 2D state space. Polar coordinates were transformed from the cartesian coordinate of the first 2 principal components. The color of each trajectory indicates the duration (cool to warm reflecting short to long words), and transparency indicates relative elapsed time from word onset to offset. D. Top row: In the polar coordinate space, the radial component is correlated with duration (left), but not relative elapsed time (middle), quantified by pearson correlation (right). Each dot in the right plot indicates a time bin (for testing duration, we controlled relative elapsed time with a 10% time bin; for testing relative elapsed time, we controlled duration with a 40ms time bin). Bottom row: The polar angle is correlated with relative elapsed time (middle) but not word duration (left), quantified by rotation speed, defined as the rate at which the polar angle changes with respect to time (right). Colors correspond to bins of duration/time.

To understand whether and how time coding may occur throughout spoken word processing, we visualized the average response to words binned by duration for all word boundary electrodes. The motivation is to preserve the coding of any temporal structure that is shared across words with the same length, irrespective of specific speech content (e.g., phonetic), by taking the average neural response over these trials (**Fig. 3B**, top).

To visualize and quantify this temporal structure, we applied dimensionality reduction to characterize the state-space trajectory of population cortical activity. For example, recent studies have provided compelling evidence that low-dimensional activity in the motor cortex can be largely explained by inherent dynamical interactions (Churchland et al., 2010, 2012; Michaels et al., 2016). In cognitive tasks, medial frontal neurons demonstrate firing rate profiles that are temporally scaled to match the temporal expectations of events (Wang et al., 2018). Here, we investigated whether similar approaches could address the representation of time in word processing.

To understand how these neural populations collectively represent time (Buonomano & Merzenich, 1995; Cao et al., 2022; Hardy & Buonomano, 2016), we projected the binned neural response into a 2D state space through principal component analysis (PCA) (**Fig. 3B**, bottom). We observed the temporal pattern of population codes were “stretched” according to the duration of words, implying the overall movements in the neural state were delayed for longer words (**Fig. 3B**, top). Furthermore, the corresponding trajectories in the state space showed larger cycles for longer words, implying a relationship between response amplitude and word length (**Fig. 3B**, bottom). Here, we modeled each segmented unit of word processing as a circle, and transformed the first two principal components into radius and polar angle using standard definitions:

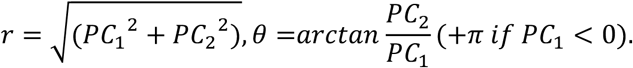

We visualized the neural trajectories in this state space across different word durations in the polar coordinate space (**Fig. 3C**). We found that regardless of word duration (from 300ms to 680ms), the neural trajectory always formed a circle that starts at the same polar angle of 48.7±10.5 degrees. Longer words traversed paths with wider radii, indicating tracking of absolute duration. Consistent with the observation in single electrodes at word boundary, this suggests that these neural populations continuously track the elapsed time from word onset. Additionally, the phase of this cycle - which corresponds to how much of the current word has been heard (elapsed time from word onset normalized by the duration of the word, which we call relative elapsed time) - resets this temporal tracking at word boundaries.

To quantify these effects, we evaluated the relation between the cyclic dynamics and temporal coding. We used the radial coordinate (**Fig. 3D top**) and the rotation speed (**Fig. 3D bottom**), i.e., the rate of change in polar angle relative to changes in time, to describe the dynamics of the cycles derived from neural data. For the temporal context of word processing, we used the absolute time (elapsed time from word onset in milliseconds) and relative elapsed time (after normalizing by word duration) to label each data sample. To understand how temporal context was reflected in these cyclic neural dynamics, we compared the pairwise correlation between these variables across different word lengths. We found that the coding of absolute duration and relative elapsed time could be distinctly characterized by the radial coordinate and polar angle, respectively. Specifically, the radial coordinate consistently reflects the absolute time (r=0.40±0.33; one-sample t-test p<0.01), not relative elapsed time (r=-0.12±0.46; one-sample t-test p=0.50) (**Fig. 3D top**). On the other hand, the rotation speed remained constant across word lengths when assessed with relative elapsed time (5.72±0.65 rad/cycle), but not with absolute time (6.35±2.01 rad/500ms; F-test comparing the variance, F stat = 9.60, p<0.001), suggesting the phase of these rotations faithfully tracks the relative elapsed time throughout the word (**Fig. 3D bottom**).

Together, these results demonstrate that as words unfold in continuous speech, neural populations precisely track how much of each word has been heard. This allows for the cycle of acoustic-phonetic, prosodic, and lexical feature encoding (**Fig. 2**) to operate consistently in the face of massive variability in both the content and duration of words. Importantly, our findings suggest that these neural populations adapt word encoding to accommodate varying durations during online word processing.

### Word-level representations emerge in a self-supervised learning model of speech

Word perception is natural for humans, yet has been a traditionally difficult computational challenge for automatic speech recognition (ASR) systems given the lack of reliable acoustic cues as previously explained. Recent deep learning approaches have achieved human-like performance, and new comparative studies have revealed similar representations for phonological processing between biological and artificial neural networks (Hsu et al., 2021; Li et al., 2023). In this study, we specifically explored an advanced self-supervised speech learning model (HuBert) (Hsu et al., 2021). This model is trained to predict the “pseudo-labels” of masked audio inputs based on unsupervised clustering, instead of being explicitly trained to identify words in continuous speech. However, its large number of parameters distributed over a deep transformer architecture allows it to discover this crucial unit (Kamper, 2023; Pasad et al., 2023; Peng & Harwath, 2022). Here, we ask where in the model these word unit representations emerge, and how the dynamics that generate these representations correspond to neural data from the human brain.

To test whether the model learns to identify word boundaries through self-supervised learning, we applied the same decoding analysis (binary logistic regression classifier with L2 regularization) to the artificial neurons in HuBert (30 PCs from 1024 units in each transformer layer; number of layers = 25). We then compared the results with those obtained from real neural populations (**Fig. 1**). We found that all transformer layers showed above-chance word boundary decoding, with increasing performance toward deeper layers (max at layer 21; AUC=0.95; **Fig. 4A**), which was slightly higher than the decoding performance from all neural populations (orange line, AUC=0.91; grouped permutation test for paired AUCs, p=0.0015). Except for the earliest layers of the model, word boundary decoding was significantly better than a decoder trained on the acoustic spectrogram (AUC=0.83; grouped permutation test for paired AUCs, p<0.001 for layers after layer 8) or envelope (AUC=0.72; grouped permutation test for paired AUCs, p<0.001 for all layers). These results suggest that word boundary effects observed in STG neural populations may reflect similar processes as the mid-deep layers of a high performance self-supervised speech recognition model.

**Fig. 4:**
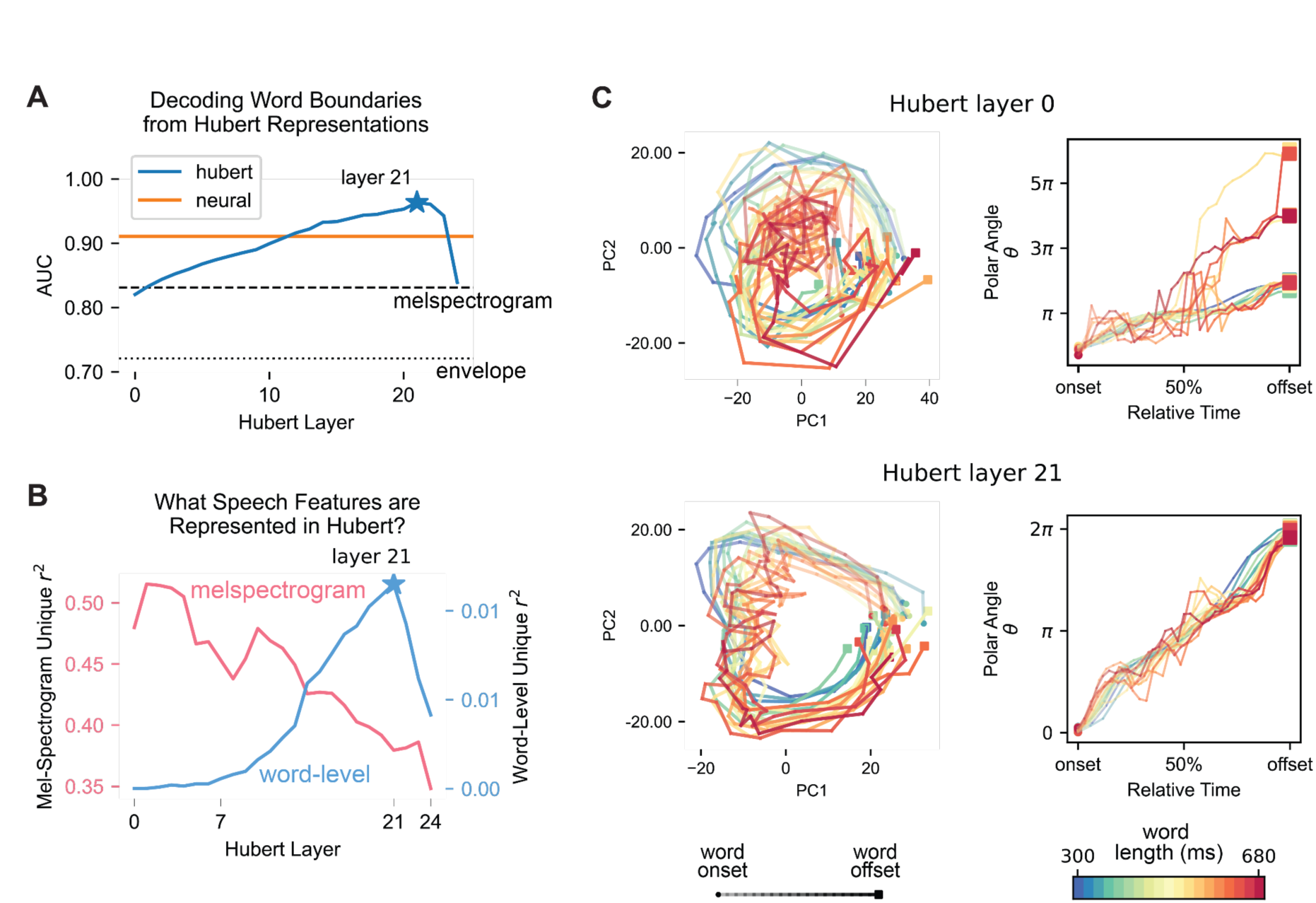
A self-supervised, deep learning speech recognition model mirrors the word boundary behavior, feature representation, and temporal dynamics of human STG. a. Word boundary decoding performance is highest at mid-deep layers of a transformer-based self-supervised learning speech model (HuBert). Specifically, at layer 21, word boundaries are decodable with near perfect accuracy (AUC = 0.95), comparable to the decoder trained on all neural populations (AUC = 0.91, orange line), and significantly higher than a decoder trained with the acoustic spectrogram (AUC = 0.83, thick dashed line) and envelope (AUC = 0.72, thin dashed line). b. As information progresses through the layers of the model, activations explained by word-level features emerge in deeper layers, while acoustic spectrogram features moderately decrease, though are still dominantly encoded throughout all layers (compare left and right y-axes). c. Hubert layer 21 shows consistent cyclical word dynamics in contrast to HuBert layer 0. Left: The trajectory of the state space representations of artificial neurons color-coded by words with different length (cooler colors for shorter words and warmer colors for longer words). Right: relation between polar angle and relative time across words with different length.

To understand what speech information encoded in the deeper layers contributes to the model’s word extraction behavior, we used TRF encoding analysis to probe the feature representations in these artificial neurons. Specifically, we found that acoustic-phonetic features were represented throughout all model layers, most strongly in lower to middle layers. In contrast, word-level features (see Methods) gradually emerged from middle layers and peaked at layer 21 (**Fig. 4B, Fig. S3**). The alignment between word boundary decoding performance and word-level feature encoding across HuBert layers (pearson correlation r=0.87, correlation between blue curves in **Fig. 4A** and **Fig. 4B**) may suggest that the HuBert model also leverages lexical information to identify word forms, akin to the mechanism in human speech cortex (**Fig. 2**). Similar to the overlapping and integrative representations we observed in STG (**Fig. 2E**), even in layer 21 where the word-level feature encoding peaked, the acoustic spectrogram still explained the major portion (38.0%) of total variance in the model representation.

These findings suggest that the speech model was able to learn the boundaries and features of words without explicit supervision. We then asked whether it shares a similar computational mechanism to the human STG to track temporal dynamics during word processing. Using the same strategy for assessing temporal context in neural dynamics (**Fig. 3**), we applied the state space analysis to each layer in the model (**Fig. 4C**). Intriguingly, we found that the trajectory for long words (600ms) did not form into a cycle (**Fig. 4C, top**), which suggested that at this stage, HuBert has not yet extracted the word forms since the model could not consistently track the temporal context of words with varying length. This was consistent with the fact that the initial layers lack the necessary information for precise word units.

In contrast, in deeper HuBert layers, each word trajectory was untangled into a single cycle. This trend applied to all words, where we found the geometry of representations from word onset to word offset tended to be scattered in the input layer, but formed a circle in layer 21, tracking the progress of word processing from onset to offset, regardless of the word length (**Fig. 4C, bottom**). Similar to STG neural populations, we found that the phase of each cycle tracked relative time (**Fig. 4C, right**; rotation speed = 5.57±0.41 rad/cycle for layer 21 and 8.15±5.15 for layer 0; F-test comparing the variance, F stat=157.30, p<0.0001). In summary, the learned representations in HuBert exhibited a cyclic pattern in deeper layers when word forms can be effectively extracted, akin to how neural populations in the brain track temporal context during spoken word processing.

Collectively, our observations revealed a strong alignment between a self-supervised speech learning model and the human brain. Specifically, similar to human STG, word encoding in HuBert is distributed across layers and co-encoded with spectrotemporal speech features. Most interestingly, we identified consistent cyclic word dynamics in the deeper layers of HuBert, a phenomenon absent in the lower layers of the model. This indicates that word units are extracted by the model without explicit learning of linguistic knowledge. Furthermore, the artificial neurons demonstrate the same rotation speed regardless of word length, implying an encoding of the relative time within HuBert neurons. Together, these results imply that the self-supervised learning model shares a similar computational mechanism akin to that of the human STG when processing individual words in natural speech.

### Dynamic alignment of cortical responses to perception of word units

The results thus far demonstrate that neural word form encoding in natural speech relies on the processing of multiple acoustic and linguistic cues in STG, aligned to word boundaries. A final question we sought to address is how word form encoding relates to the listener’s perception. Even under ideal listening conditions, there are often multiple possible interpretations of speech input that lead to different word percepts; this phenomenon is the basis for so-called “mondegreens”, which are commonly mis-heard phrases or song lyrics (Wright 1954). Thus, we hypothesized that neural encoding of words is affected by the perceptual experience of the listener even when the acoustic and linguistic properties of the speech input are fixed.

To understand how STG neural population activity reflects listeners’ perception of words in the face of ambiguous input, we developed a bistable speech stimulus in which the same acoustic speech stream could be perceived as multiple distinct words depending on where listeners identify the word boundary (Warren 1961). In the first experiment, three participants listened to streams of syllables that were repeated approximately 1000 times without gaps (**Fig. 5a**). These sounds were generated to allow listeners to interpret the same sound as either the first or second syllable in a word (e.g., “turbo” /tɛrbo/ vs “boater” /botɛr/). Since the acoustic input is the same, these stimuli provide a highly controlled way of evoking purely internally-driven word form perception.

**Figure 5:**
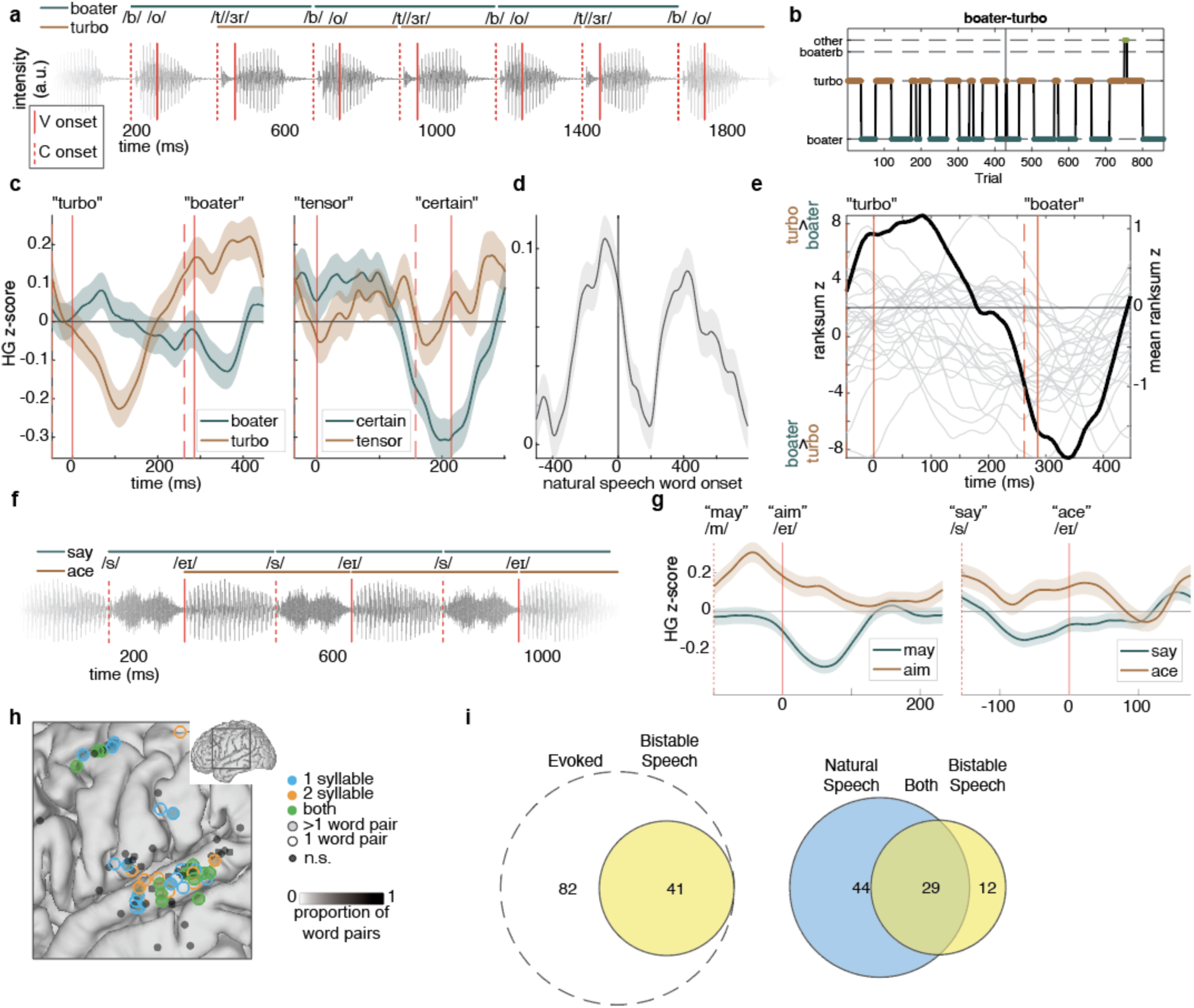
Word encoding in a bistable speech stimulus reflects listeners’ subjective perception. **a.** Example stimulus excerpt for the bistable sequence that can be heard as “turbo”/“boater”. Solid red lines mark vowel onsets, dashed red lines mark consonant onsets. **b.** Trial-by-trial perceptual reports from one representative participant for “turbo/boater”. **c.** Example electrode responses to “turbo/boater” (left) and “tensor/certain” (right), showing distinct changes in HGA aligned to perceived word onset. **d.** Word-boundary aligned response from natural speech for the same electrode in c. **e.** All electrodes with significant differences in HGA between “turbo” and “boater”. Left y-axis: individual electrodes (gray lines); Right y-axis: mean across electrodes (black line). Values reflect the test statistic from a ranksum test, with positive values reflecting turbo>boater and negative values reflecting boater>turbo. **f.** Example stimulus excerpt for the bistable sequence that can be heard as “say/ace”. **g.** Example electrode (same as in panel c-d) showing different responses to percepts for “may/aim” (left) and “say/ace” (right). **h.** Electrodes with significant perceptual effects for 1-syllable (blue), 2-syllable (orange), or both (green) types of stimuli, primarily in STG (black indicates evoked response with no perceptual effect). **i.** Half of electrodes with evoked responses to bistable stimuli show perceptual effects (left). Most electrodes with bistable perceptual effects also show significant word boundary effects in natural speech (right). Only participants with overlapping data in both tasks are included.

Participants held down buttons with either the left or right hand to indicate which word they were perceiving at each moment (a single perceived word is defined as a trial). They tended to hear the same word multiple times in a row, switching among them at variable intervals (**Fig. 5b**). Across all participants and stimuli, the two intended word percepts were the most common, while other non-word percepts (e.g., “boaterb”) were rare (**Fig. S4a**).

Next, we asked whether neural populations in STG reflect the specific word listeners hear on a given trial, despite identical acoustic input across conditions. We identified electrodes that had evoked HGA responses to the sounds in the speech stream across trials (>3 SD difference from within-trial mean across time, two-tailed). Of the 267 electrodes with significant evoked responses to these bistable stimuli, 45 (16.8%) had significantly different responses to the two perceived words (ranksum test with temporal cluster permutation, p<0.05). For example, an electrode over left STG showed a difference in the amplitude of the evoked response to the perceived words “turbo” and “boater” (**Fig. 5c**, left; ranksum test with temporal cluster permutation, p<0.05). The same electrode showed a difference for another bistable stimulus on trials perceived as “tensor” vs “certain” (**Fig. 5c**, right). The differences between word percepts were strongest within ∼100ms following perceived word onset, and were characterized by a decrease in HGA that was more pronounced for the perceived word. Notably, this same electrode was characterized as a word boundary electrode in the natural speech task (**Fig. 1**), and exhibited the same HGA decrease immediately following word onset in that task (**Fig. 5d**). These results demonstrate that neural populations in STG that track word boundaries in natural speech reflect listeners’ specific word perception on each trial.

To evaluate word boundary encoding across participants and electrodes, we examined the magnitude of the difference between word percepts across time. The time course of the test statistic (ranksum z-value) showed that the maximum difference across electrodes occurred ∼100-150ms after the perceived word boundary (**Fig. 5e**). Together, these results suggest that the human brain rapidly encodes perceived word forms in transient changes in neural activity surrounding the perceived word boundaries.

To establish that STG definitively reflects perceptual word form encoding regardless of phonological content or length, we created another set of stimuli where different words could be perceived depending on the perceived order of two phonemes within a single syllable, for example the consonant-vowel (CV) sequence, “say” (/seɪ/) versus the vowel-consonant (VC) sequence, “ace” (/eɪs/) (**Fig. 5f**). As with the 2-syllable stimuli, participants alternated primarily between the two word percepts, while non-word CVC (e.g., “sace” [/seɪs/]) and other percepts were rare (**Fig. S4b**). Neural populations in STG differentiated between different percepts on individual trials, demonstrating perceptually-driven word form encoding for even short phonological sequences (**Fig. 5g**; ranksum test with temporal cluster permutation, p<0.05; see **Fig. S4c** for quantification of perceptual word form encoding within electrode and across multiple stimuli). These results demonstrate that STG is sensitive to perceived word boundaries for single syllable words where percepts differ only in the order of the consonant and vowel.

Finally, we asked to what extent electrodes exhibiting perceptual effects in the bistable task overlapped with word boundary electrodes identified in natural speech. First, we observed that the majority of electrodes with perceptual effects were found in the mid-STG (**Fig. 5h**). Furthermore, perceptual effects in these electrodes were generally independent of word length (1- or 2-syllable) or the phonological units that define the boundary (phonemes or syllables), suggesting a general word form encoding mechanism within this region. Second, we found that out of the 82 speech-responsive electrodes that were recorded in both the bistable and natural speech tasks, 41 (50.0%) had significant effects in the bistable task (**Fig. 5i**, left). Out of these electrodes, 29 (70.7%) also showed word boundary effects in natural speech (**Fig. 5i**, right). Thus, a subset of neural populations that encode word forms in natural speech are directly influenced by trial-by-trial perception, which is driven by factors beyond just the bottom-up acoustic speech input.

## Discussion

A fundamental challenge in understanding spoken language processing lies in uncovering how the brain represents words as discrete linguistic units. Word encoding requires the integration of diverse acoustic and linguistic cues—acoustic-phonetic elements, prosody, and lexical information—over hundreds of milliseconds to form a coherent auditory word form corresponding to a holistic lexical item.

Our findings challenge traditional models that posit word form representation occurs in separate brain regions and at distinct processing stages from sub-lexical encoding. Instead, we show that neural populations in the mid-superior temporal gyrus (STG) exhibit continuous, dynamic encoding of phonetic, prosodic, and lexical features in a processing cycle, which is “reset” at word boundaries. This dynamic processing positions the mid-STG as a critical integrative hub for spoken language, expanding its role beyond basic spectrotemporal analysis (Hickok & Poeppel, 2007). Indeed, previous studies have demonstrated neural populations in STG that are specialized for speech processing over other sound inputs, such as music and environmental sounds (Norman-Haignere et al., 2015; Sankaran et al., 2024). While there is certainly a role for other regions within the broader cortical networks associated with language (Fedorenko & Thompson-Schill, 2014; Fernandino & Binder, 2024), the high spatiotemporal resolution of ECoG recordings here reveals the strikingly diverse set of operations that exist within the STG, including at the level of words.

One of the most striking findings is that STG neural activity identifies word boundaries and maintains a consistent temporal tracking across all words with variable speech contents. The stronger neural suppression after longer words may reflect a “washout” of the temporal integration window, whose size depends on the amount of phonological information temporarily stored in working memory to allow encoding of the whole word form. In addition, the neural dynamics track the relative time throughout the word, implying that these neural populations are able to represent the temporal context in real time, normalized by duration. This normalization parallels mechanisms in visual object recognition, where the brain computes size-invariant representations that allow diverse sensory inputs to map onto stable stored representations (e.g., face recognition across changes in angle or lighting) (DiCarlo et al. 2012). This enables an internal mechanism to bind phonological sequences over time into duration-invariant representations of word form.

We observed similar word-level dynamics in the deeper layers of a self-supervised speech recognition model (HuBert), despite it not being explicitly trained to recognize words. Compared to human STG, we found similar behaviors, representations, and dynamics when extracting individual words from continuous speech. The duration-invariant tracking of relative time within words was observed only in the higher layers of the transformer model after several non-linear transformations of the spectrotemporal information. This is likely enabled by the model’s positional encoding - the key mechanism in transformer models (Vaswani et al. 2017) that handles sequence order, which acts as an “internal clock” and is linearly added to the representation of speech content. As a result, the representation of phonological information in HuBert becomes sequence-dependent, allowing for the dynamic encoding of word forms. Taken together, these findings suggest that, like the STG, HuBert solves the word extraction task through a dynamic and integrative function acquired via passive listening. These results provide novel insights into the mechanistic-level similarity between the human brain and AI speech models, beyond what earlier studies have shown at the representation level (Caucheteux et al., 2023; Goldstein et al., 2022; Li et al., 2023).

A key demonstration of the functional relevance of these neural dynamics comes from our bistable word perception task. Here, STG activity is aligned with subjective word perception on a trial-by-trial basis, reinforcing the idea that neural activity reflects not only acoustic input but also perceptual interpretation. This tight coupling between physiology and perception is essential, given the inherent ambiguities in natural speech. Spoken language often involves non-linear mappings between acoustic signals and linguistic categories, requiring listeners to deploy rapid, parallel computations to derive meaning (Fox et al., 2020; Leonard et al., 2016).

Collectively, our findings support a dynamic, integrative model of auditory word form processing. Rather than relying on discrete stages or regions for different components of word processing, the STG encodes word forms through continuous population dynamics that integrate acoustic and linguistic information in real-time. This framework bridges well-characterized phonological representations in the auditory cortex with higher-order linguistic representations that enable meaningful speech communication. By revealing the dynamic neural code underlying word forms, this study advances our understanding of how the brain transforms continuous speech into meaningful linguistic units. Future work could extend this model to explore how these dynamics interact with broader language networks and contribute to speech production and comprehension in more complex linguistic contexts.

## Methods

### Participants

The University of California, San Francisco Institutional Review Board approved all procedures, and all patients provided written informed consent to participate.

Sixteen participants were implanted with high-density ECoG grids (256-channel, 16x16, 4-mm distance) as part of their treatment for intractable epilepsy. Grid placement was determined by clinical considerations (12 left hemisphere, 4 right hemisphere). Electrodes were localized by aligning pre-implantation MRI and post-implantation CT scans (see Fig. S1 for the electrode coverage).

Seven participants (4F/3M) performed the bistable perception task, and were implanted with the same high-density ECoG grids as above. Of these seven, two also listened to the naturally spoken narratives, allowing for direct comparison across tasks.

### Stimuli and Tasks

Sixteen participants listened to 10 short (181s ± 29s) spoken narratives from the Boston University Radio News Corpus (BURSC) (Ostendorf et al., 1995). BURSC was used because it contains natural continuous speech, and speech samples were carefully transcribed with phonetic, prosodic, and word annotation by linguists.

The bistable perception experiment was created following an earlier approach known as verbal transformation (Warren 1961). A male speaker was recorded saying a set of six words (“turbo”, “tensor”, “say”, “day”, “may”, and “zoo”). Each sound file was trimmed to identify start and end samples where the amplitude was 0dB, and then that trimmed section was concatenated between 325-540 times to produce a continuously repeating stimulus lasting ∼3 minutes. Each stimulus was repeated twice, for a total of 650-1080 trials, with each trial being the onset of a word in the continuous stimulus. Participants were instructed to hold down buttons with each hand to indicate their continuous perception of each stimulus as a repeating word. For example, they would hold down the left hand button to indicate when they perceived the sound as “say”, the right hand button for “ace”, both buttons for “sace”, and neither button for “other”. Across the two repetitions of each stimulus, the mapping of the two primary word percepts was counterbalanced across hands.

### ECoG data pre-processing

Electrophysiological recordings were acquired at a sampling rate of 3051.8 Hz using a multi-channel amplifier optically connected to an RZ2 digital acquisition system (Tucker-Davis, Technologies, Alachua, FL, USA).

The recorded local field potential was first notch-filtered at 60Hz and its harmonics to remove line noise and then bandpass filtered in the high gamma range (70-150Hz) using the Hilbert Transform at 8 logarithmically-spaced center frequency bands within this range. We then took the average amplitude across these eight bands. This high gamma activity (HGA) was further downsampled to 100Hz and z-scored across the entire recording block for the following analyses.

### Decoding word boundaries from single-electrode HGA

To assess whether neural populations exhibited distinguishable temporal dynamics to word boundaries compared to within-word syllable boundaries, we conducted a set of decoding analyses for individual electrodes. For simplicity and interpretability, we trained binary logistic regression classifiers with L2 regularization from the neural activity around unit boundaries.

First, we aligned the neural activity to all word (n=1587) and within-word syllable (n=2967) boundaries. Note that in this paper, we only including multisyllabic words, since word and syllable boundaries are confounded in monosyllabic words, and since multisyllabic words provide longer windows to allow us to evaluate multiple peaks in the evoked neural response.

Then, we used the preprocessed HGA within a temporal window around these boundaries as the predictors for the decoder. Specifically, when optimizing the decoders we utilized a sliding window approach and tried three different window sizes (window = 100ms, step = 10ms; window size = 500ms, step = 50ms; window = 1200ms, a single step). The sliding window analysis started from 500ms before the unit boundaries and extended to 700ms after the unit boundaries. The 1200ms window, which covered the entire search range, was referred to as the “full window”.

Unless specified otherwise, we primarily present the results obtained from the 500ms window size. We opted for this window size as further enlarging it to the full size did not yield substantial improvements in accuracy.

Since the preprocessed HGA was sampled at 100Hz, we derived 50 temporal predictors from the 500ms window, denoted as the vector 𝑋*_i_* where 𝑖 refers to a specific sample. The corresponding target 𝑦*_i_* indicates the underlying class label, distinguishing between a word boundary (𝑦_i_ = 1) and a syllable boundary (𝑦*_i_* = 0).

To fit the model, we solved the following optimization problem:

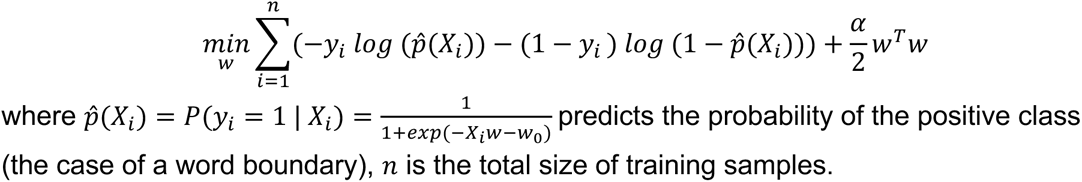

We implemented the L2 regularized logistic regression with the Python package `scikit-learn` (C=0.01; random_state=0; max_iter=100). In our approach, we randomly sampled 500 instances from both classes for training, and reserved the remaining data for testing. The decoder’s performance was assessed by the AUC score (area under the receiver operating characteristic curve, or ROC curve), which provides a single number (range from 0 to 1; chance level=0.5) that accounts for both the sensitivity and specificity of a binary classifier. This procedure was repeated 10 times, with each iteration involving a different randomization for the train-test split. We then evaluated the performance of the decoder by averaging the results obtained on the testing set across these iterations, separately for each electrode.

To evaluate the statistical significance of these neural decoders, we utilized a permutation test by conducting the training process with randomly shuffled class labels (𝑦_i_). We then compared the decoder’s performance when trained with correct labels against that when trained with shuffled labels. Specifically, we calculated the p-value through an independent one-sided t-test by comparing the AUC values, which were derived from 10 different train-test splits for both the original and shuffled conditions. Electrodes exhibiting significantly higher decoding performance (p<0.001) were considered as word boundary electrodes.

To evaluate the statistical significance between the decoding performance from two decoders, we applied a group permutation test to paired AUCs. The null hypothesis was that there was no difference in AUC between the two decoders. To generate the null distribution of AUC differences under the null hypothesis, we combined the predicted scores from both decoders into a single array and created corresponding group labels indicating the source decoder for each score. We then performed 10,000 permutations, where in each permutation, the group labels were randomly shuffled to break any association between the predicted scores and the decoders. For each permutation, the shuffled group labels were used to reassign the prediction scores into two groups. The AUCs for the permuted groups were calculated with the true labels. The p-value was determined by calculating the proportion of the permutation distribution where the permuted difference in AUCs was greater than or equal to the observed difference.

For the bistable perception task, the HGA ECoG data were epoched relative to either the consonant (for the one-syllable stimuli) or the first consonant of one of the possible word percepts (for the two-syllable stimuli). To identify electrodes with significant differences between trials perceived as different words, we used a cluster-based permutation approach (Maris and Oostenveld 2007) to detect timepoints where temporal clusters had significant z-values for a ranksum test between conditions at p<0.05.

### Characterizing peaks from neural activity at word boundaries

To explore the temporal dynamics of neural activity at word boundaries, we categorized three potential types of peaks. These categories were defined based on common patterns observed in the word boundary electrodes: a trough immediately following the word boundary, a positive peak preceding the trough, and another positive peak following the trough.

To investigate the properties of these peaks, particularly their timing, we used the following steps to detect if a word boundary electrode exhibits one or more of these peaks. For each electrode, we first calculated the word event-related potentials (ERPs) by averaging the neural responses aligned to word boundaries (weighted by the decoding score), which resulted in a single time series encompassing a time window from 500ms before the word boundary to 800ms after the word boundary. Then, we detected if there is a significant trough (prominence=0.05, width=50ms) in the word ERP using a peak-finding algorithm. If such a trough was identified, we further searched for significant positive peaks (prominence=0.08, width=50ms) before or after the trough.

### Decoding word boundaries from populations of electrodes

We further conducted a set of decoding analyses for electrode populations. First, for each subject, we identified all the electrodes that demonstrated a significant word boundary response (as identified by the decoding performance; permutation test p<0.001), and the median number of boundary electrodes per subject was 21 (minimum=6, maximum=42). Then, for each word boundary electrode, we cropped a 500ms window around the negative boundary peak. These temporal responses were concatenated as the input feature for the neural population decoder defined separately for individual subjects.

As with single electrodes, we trained binary logistic regression classifiers with L2 regularization from neural response around unit boundaries. For each decoder, we randomly sampled 500 instances from both classes for training, and reserving the remaining data for the testing purpose. This procedure was repeated for 10 times, with each iteration involving a different randomization for the train-test split. We then evaluated the performance of the neural population decoder by averaging the results obtained on the testing set across these iterations, separately for each subject.

### Extracting speech features from stimuli

We extracted speech and language features from the BURSC stimuli for training the encoding model. These features can be categorized into several groups, including spectrogram features, envelope features, pitch features, phonetic features, phoneme statistics features, prosodic features, boundary features, lexical features, part-of-speech features, and large language model features.

### Acoustic-phonetic features

1. Spectrogram features: We used the Python library `librosa` to extract mel-spectrogram features from the BURSC stimuli. The audio wave for each stimulus (sampled at 16,000 Hz) was first transformed into its power spectrogram (FFT window = 100ms, step = 10ms), and then mapped onto 16 mel-scale frequency bins (0 ∼ 8,000 Hz) to align with human perception.
2. Continuous envelope magnitude: We applied the Hilbert transform to get the amplitude envelope of the original audio signal. Next, we downsampled it to 100Hz to match the sampling rate of the high gamma signal, and a third-order Butterworth low-pass filter was applied for temporal smoothing. Finally, we normalized the envelope values between 0 and 1.
3. Envelope peak rate: To obtain the peaks of the envelope, we used a peak-finding algorithm to identify timepoints where the magnitude was 10% of the loudest point over the whole stimulus. To obtain the peak rate feature (Oganian & Chang, 2019), we applied the same peak-finding approach to the first-order temporal derivative of the normalized continuous envelope (threshold = 0.1).

### Prosodic features

1. Vowel lengths and syllable lengths: were defined as the elapsed time (in milliseconds) between the onset and offset of the unit.
2. Pause duration: the gap (in milliseconds) at word boundaries (equals to 0 for 91.9% samples in this natural speech stimuli)
3. Pitch: We extracted four distinct pitch features to characterize the speaker’s fundamental frequency (F0). We applied the algorithm implemented in the Python library `CREPE` to estimate the pitch contour from the original audio signals, which was then downsampled 100 Hz, smoothed using a 100ms window, and normalized between 0 and 1 within each stimulus to obtain a measure of speaker-normalized (relative) pitch (Tang et al., 2017). We also extracted three discrete features to capture the dynamics in pitch. We identified the peak pitch as the highest pitch from the normalized pitch contour within each ∼30 second block of speech. We then calculated the pitch-up and pitch-down from the first-order temporal derivative of the normalized pitch contour, which characterized the increasing change (positive peak) and decreasing change (trough) respectively.

### Sequence statistics

1. Biphone surprisal: we estimated biphone surprisal from the SUBTLEX-US corpus , similar to prior work (Brodbeck et al. 2022; Leonard et al. 2015). We first concatenated the phoneme sequence (across word boundaries) but kept each sentence separate. We then estimated the distributions and conditional probabilities based on the first phoneme identity in each biphone sequence < 𝐴𝐵 >:

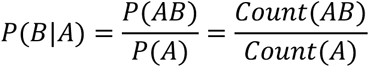

Then, we defined the biphone surprisal feature as − 𝑙𝑜𝑔_2_ (𝑃(𝐵|𝐴)).

### Lexical features

To derive these features for the BURSC stimuli, we utilized the vocabulary set of commonly spoken words (as indicated by the column CobS) from the CELEX database (Baayen et al. 1996), which provides the spoken frequency of 16,276 lexical items.

1. Word duration: Word duration was defined as the absolute time (in milliseconds) elapsed from word onset to word offset.
2. Word frequency: Word frequency was directly adopted from the CELEX database (spoken word frequency: CobS) and transformed to the logarithmic scale for the following analyses.
3. Cohort density: For each word in the BURSC stimuli, we counted its neighbor word phoneme by phoneme and transformed the results to the logarithmic scale. We adopted the CMUdict (the CMU Pronouncing Dictionary) to get the pronunciation of each word as a sequence of phonemes.
4. Uniqueness point: The uniqueness point was defined as the offset of the phoneme where the stems of the cohort words became incompatible.

### Partial correlation analyses

We used partial correlation to estimate the relationship between HGA and the speech information. For the k-th word boundary trial, we collected all the speech features, denoted as {𝑥_i_^k^ | 𝑖 ∈ 𝑆}, S being the feature set as mentioned in the previous section. For the n-th electrode, we investigated the HGA from -450ms before the boundary to 650ms after the boundary, denoted as 𝑦_n_^k^(𝑡). We grouped trials into 5-sample bins by sorting the neural features, to increase the signal-to-noise ratio for correlation analyses.

For each electrode, we did this correlation analyses for each time point in 𝑦*_n_*(𝑡), which resulted in a time series of correlation coefficient around the word boundary. These analyses were implemented using the inverse covariance matrix (Kim 2015) through the python package `pingouin`(Vallat 2018).

### Decoding word duration from boundary decrease

To quantify how accurately the boundary decrease encodes the information about word duration, we applied decoding analyses at both single-electrode and population levels. For all electrodes with a prominent boundary decrease (n = 319), we collected the HGA at the negative peak for all words that were not at the beginning or end of the sentence (n trials = 4419). We trained a linear regression model with L2 regularization to predict word duration from this response. For population-level decoding, we concatenated the negative peak response across all electrodes as the multivariate predictor. The regularization parameter was optimized on a held-out evaluation dataset (10-fold cross-validation). We further applied a permutation test by shuffling the word duration labels. The statistical significance was evaluated by the Mann-Whitney-Wilcoxon test (two-sided).

### State space analyses

To assess the temporal dynamics of neural population during spoken word processing, we first collected all the boundary electrodes to form a “meta-subject”. We then extracted the high gamma amplitude from word onset to word offset for each individual word (including the monosyllabic words, n=5280 for the whole stimuli dataset) and each boundary electrode (n=321).

Here, we hypothesize the activation of a neuron 𝑁 could be viewed as a function of these two variables:

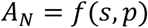

where 𝑠 is a variable describing the speech content (e.g., a phonetic feature), and 𝑝 is a variable describing the position within a word (e.g., the elapsed time from word onset).

The concept of content code is defined as an evoked neural response linked to a particular speech feature (e.g., sound /s/), regardless of its position within the word (“save” or “face”). In contrast, the concept of position code is defined as an evoked neural response linked to the “position” (which can be relative time or absolute time) within a word, regardless of the speech content present at that moment. In a standard temporal respective field (TRF) model, we usually assume 𝐴*_N_* primarily depends on 𝑥 but not 𝑝, so we could estimate the activation function by taking the expectation over 𝑝:

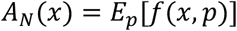

Conversely, as now we want to investigate the potential neural encoding of the position code, we estimate the activation function by taking the expectation over 𝑥:

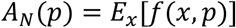

Through this analysis, we aim to get insights into how the neural representations vary over the relative or absolute time within the processing of a word segment. Note that here we explicitly tested the temporal coding for word processing, so we reset 𝑝 to 0 after each word boundary.

To achieve this, we estimated 𝐴*_N_*(𝑝) by grouping word trials according to their length, from 300ms to 680ms, with step size=20ms. Note that we excluded the first and last words in sentences because they tend to trigger strong onset or offset responses. Groups with less than 10 trials were excluded from the following analyses. This resulted in 19 groups, and the median number of trials per group was 84. For each group, we averaged the temporal response from word onset to word offset, which resulted in a trajectory in the N dimensional populational neural space. Each time point in the trajectory could be viewed as a feature in the neural space. We obtained 941 features by collecting them from 300ms words to 680ms words. To unveil the underlying structure of this high-dimensional space, we applied principal component analysis (PCA) [Fig. 3] to projecting these features into a 2D state space. We preserved the first two principal components for the following analyses because the explained variance ratio significantly decreased in the 3rd PC (1st PC: 27.61%, 2nd PC: 14.21%, 3rd PC: 4.27%).

We then visualized how these features were organized according to word processing phase and word length in the reduced 2D state space using different color-coding strategies [Fig. 3]. The word processing phase, indicated by the transparency value, was defined as the relative time elapsed from the word onset to word offset, ranging from 0 (word onset) to 1 (word offset). The word length, represented by the color warmth, was defined as the absolute elapsed time from word onset to word offset (in milliseconds), which was consistent with the previous definitions in this paper. Note that we only used unsupervised dimension reduction methods, thus these temporal annotations were not available to the model.

### Assessing time encoding in the polar-coordinate state space

After projecting neural response into the 2D state space, we used the polar coordinates to parameterize the neural trajectory. We applied the standard transformation:

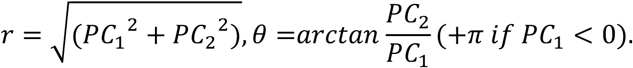

This allows us to assess how temporal information is characterized by the two parameters that define the cyclic dynamics: the radius 𝑟 (the size of the cycle) and the polar angle 𝜃 (the phase of the rotation). Specifically, for each neural trajectory in this state space, we described the absolute and relative time of individual time points. The absolute time was defined as the elapsed time in milliseconds from word onset, while the relative time was defined as the absolute time normalized by the word length (thus ranged from 0 to 1). We first assessed which temporal information the radial coordinate reflects. To quantify the relation between radial coordinate and absolute time, we first grouped individual time points according to their relative time to control for the strong correlation between these two features. We binned trials range from 10% to 100% relative time, with bin width and step both equal 10%. Within each bin, we calculated the Pearson correlation between the radius and the absolute time. Similarly, when quantifying the effect from relative time, we first binned trials by absolute time (100ms to 500ms with 40ms bin width) and then calculated the correlation between radius and relative time. We then used a one-sample t-test to evaluate if the correlation between radius and absolute time is non-zero regardless of the relative time (and vice versa).

We then assessed how temporal information affects the polar coordinate. Instead of directly using individual phase, we quantified the rotation speed inspired by the linear trends within each trajectory (Fig.3D bottom). Specifically, for words with different duration, we calculated how fast the phase changes with regard to absolute time (per 500ms) and relative time (per word, i,.e., 0% to 100%). Intuitively, this is the slope shown in Fig.3D bottom.

### Assessing speech representations in the HuBert model

We conducted a similar set of analyses on the artificial neurons extracted from a pretrained HuBert model to understand how a computational model, designed for the representation learning of human speech, achieves the capability of word processing.

First, we trained binary logistic regression classifiers with L2 regularization to distinguish between word vs. syllable boundaries for the responses in artificial neurons in HuBert. We presented the same BURSC stimuli to the HuBert model and extracted the temporal responses (sampled at 50Hz) from 1024 artificial neurons in each transformer layer (25 layers in total). To reduce the dimensionality of HuBert representations, we applied PCA to retain the first 30 principal components from 1024 artificial neurons separately for each layer. The features within a 500ms window centered at unit boundaries were used as input to the classifier. For training, we randomly sampled 500 instances from both classes, reserving the remaining data for testing. This process was repeated 10 times, and the decoding performance was quantified by the AUC on the testing data set averaged across 10 trials.

Next, we trained a TRF encoding model to assess the explainable variance in the learned representations from each HuBert layer. We used the same set of features extracted from the BURSC stimuli and resampled them at 50Hz to match the sampling rate of HuBert units. Similarly, we trained a full model and a set of reduced models to calculate the unique variance from different combinations of feature sets. Specifically, we separated features into 7 groups: mel-spectrogram, envelope, sentence onset, pitch, phonetic, phoneme statistics, word-level features (including word boundary and other word features). We then estimated the 𝑅^2^ by adding one set of feature once at a time (note that we considered mel-spectrogram as the lowest level acoustic feature):

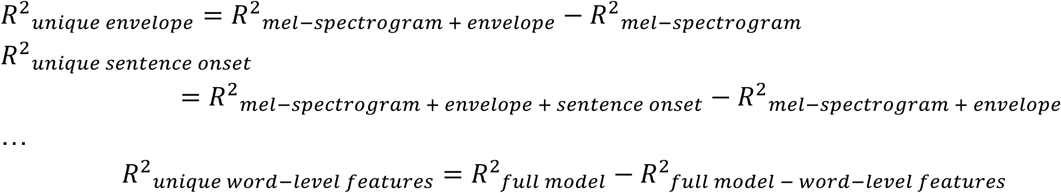

For 𝑙-th layer, the proportion of unique variance explained by a adding the 𝑘-th feature set 𝐹*_k_* was calculated by

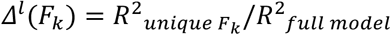

To compare how this 𝛥^l^(𝐹_k_) changed across 25 HuBert layers for different feature sets, we normalized 𝛥^l^(𝐹_k_) by 𝑚𝑎𝑥_l_𝛥^l^(𝐹_k_) for each 𝐹_k_ separately for visualization.

Lastly, we applied the state space analyses to HuBert units. For all word trials (n=5280), we extracted responses in individual HuBert units from word onset to word offset, then group them according to word length (from 300ms to 680ms, with step size=20ms). Then we applied PCA to each layer separately to retain the first 2 principal components (for layer 21, 1st PC explained 17.6% variance, 2nd PC explained 14.4% variance, 3rd PC explained 5.5% variance). We projected each time point (sampled at 50Hz) into this 2D state space and color-coded by word length (in color warmth) and word processing phase (in the transparency value) to reveal the low dimensional structure of word-level representation in each HuBert layer.

## Acknowledgements

We thank Neal Fox for assistance creating the bistable stimuli, and Emily Stephen for help with data pre-processing. This work was supported by the National Institutes of Health (grant NIDCD R01DC012379).

## Author contributions

This study was designed by Y.Z., M.K.L, and E.F.C. Funding for this research was acquired by M.K.L and E.F.C. Implantation surgeries were conducted by E.F.C. Data processing and analysis was performed by Y.Z. and M.K.L. Interpretation of the data and analysis was performed by Y.Z., M.K.L., L.G., I.B.G, and E.F.C. Y.Z. and M.K.L. wrote the first draft of the manuscript and prepared all main and supplementary figures. All authors contributed to editing and revising the manuscript.

## Competing Interests

E.F.C. is an inventor on a pending UCSF patent application (Application number: WO2022251472A1, 2022, WIPO PCT - International patent system) and patents PCT/US2020/028926, PCT/US2020/043706, and US9905239B2. EFC is co-founder of Echo Neurotechnologies, LLC. All other authors declare no competing interests.

## Code and Data Availability

All matlab code used to generate main and supplementary figures will be made available at https://github.com/YizhenZhang/word_extraction_repo.

Data to replicate all figures will be made available on request to the corresponding author and will be posted to Zenodo.

---

**Fig. S1:**
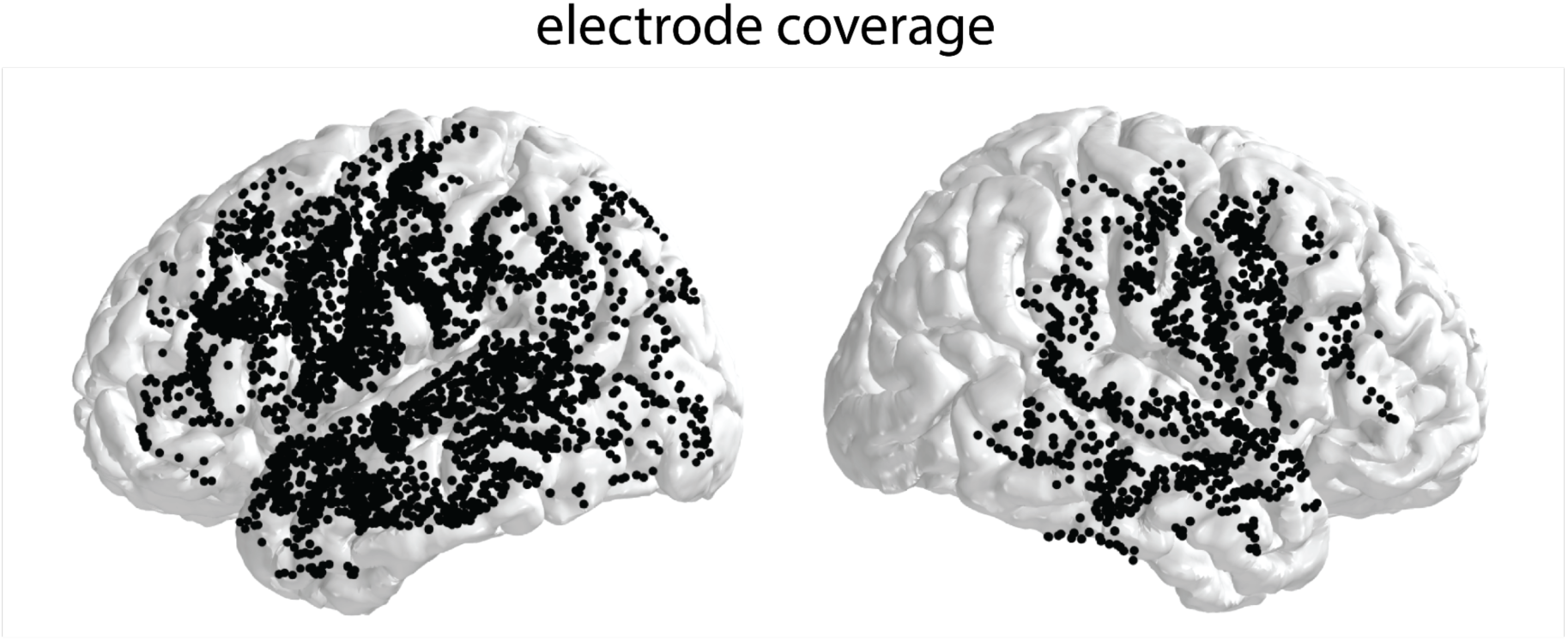
High-density electrode coverage across 16 patients (12 left, 4 right).

**Fig. S2:**
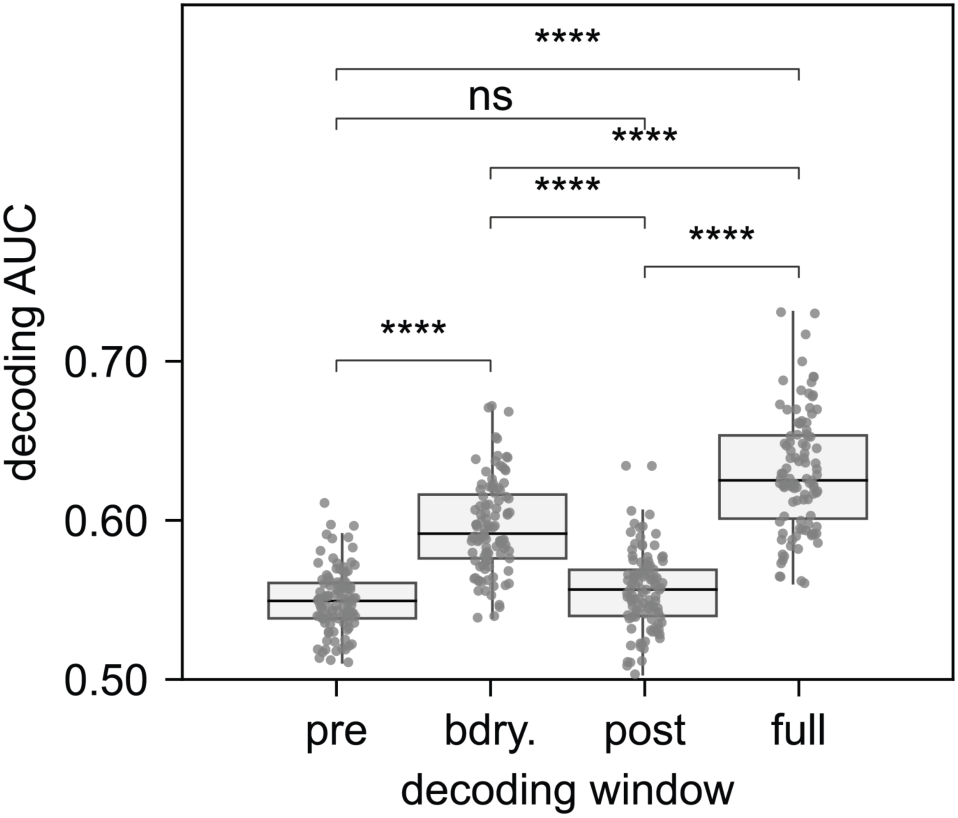
Boundary decrease is the most robust and universal neural marker for word boundaries. Among all word boundary electrodes, we observed a consistent trend that the larger decoding window (500ms before word boundaries to 700ms after word boundaries), which encompasses all peaks, consistently showed significantly superior performance compared to the narrow window (window size = 100ms) centered around single peaks (paired Wilcoxon signed-rank test, p<0.0001).

**Fig. S3:**
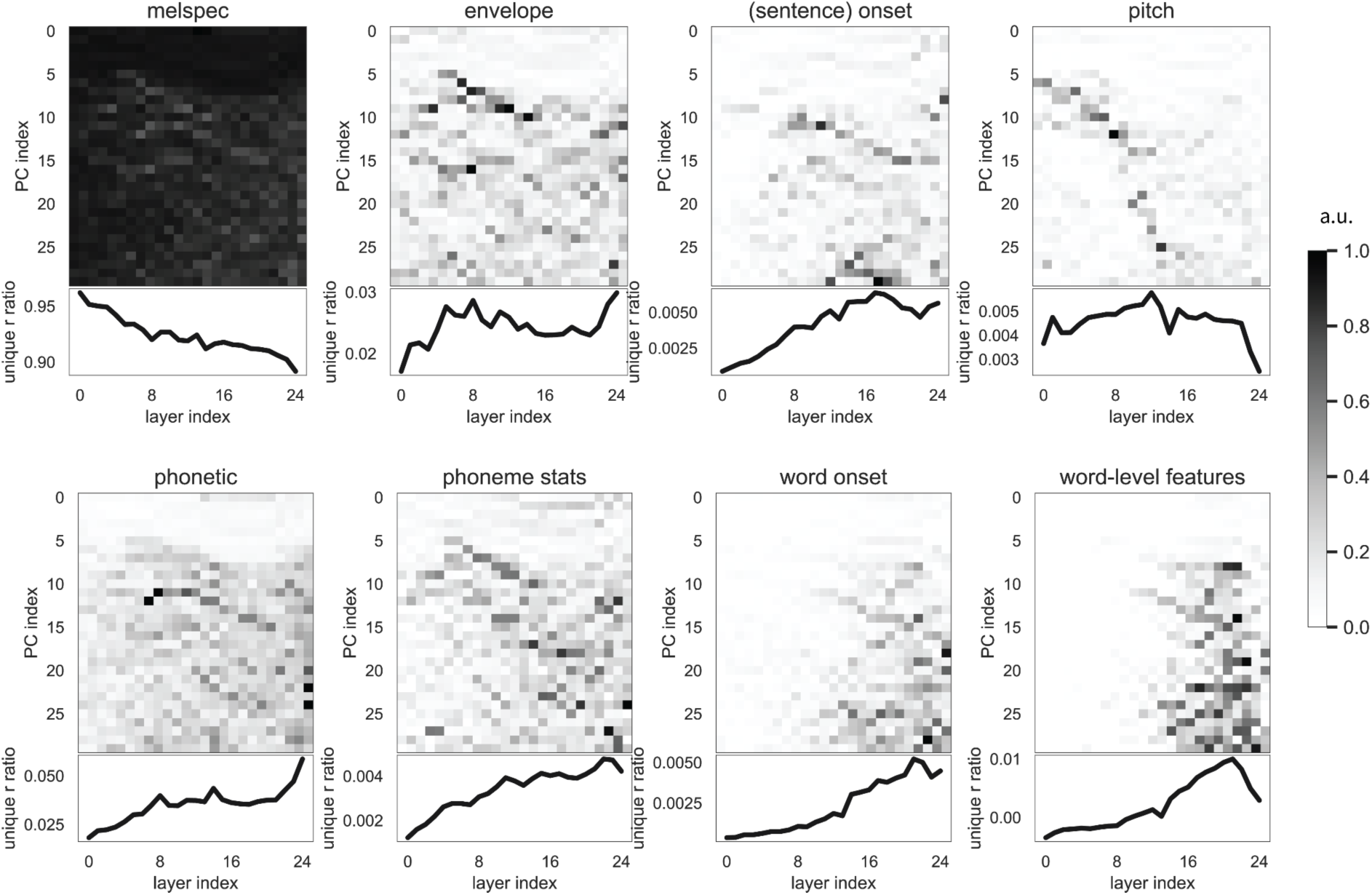
The word-level sparse information that emerges through nonlinear transformations helps improve word boundary decoding performance in higher Hubert layers. For each feature group, the 2D heatmap shows the unique variance (normalized to 1 to show the trend across layers and PCs) explained by that feature.

**Fig. S4:**
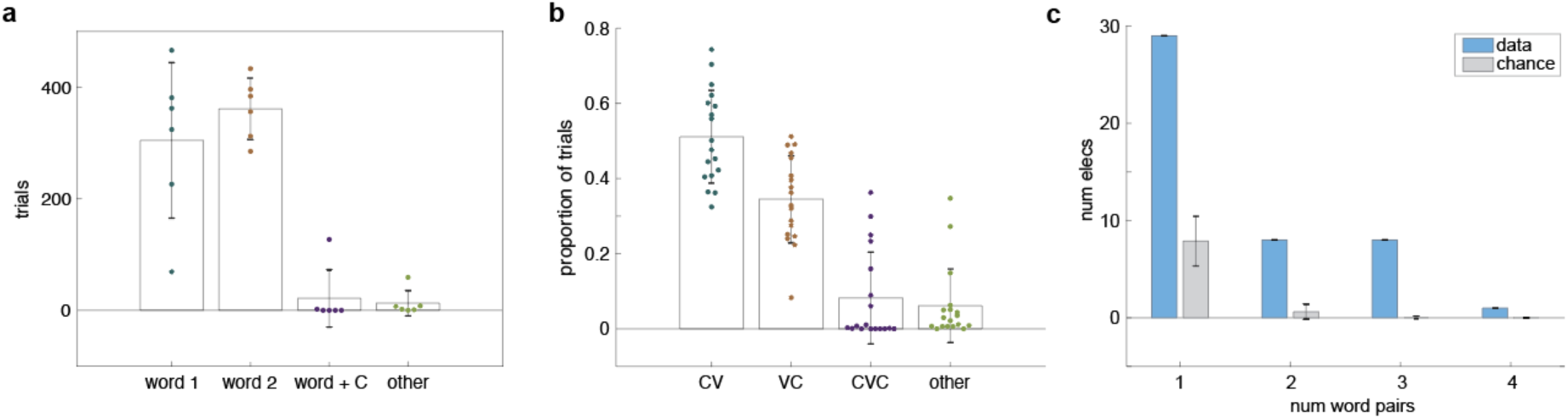
Behavioral and multi-word encoding in the bistable perceptual task. **a.** Perceptual behavior for all participants who listened to both 1- and 2-syllable stimuli (n=3). **b.** Proportion of trials perceived as consonant-vowel (CV), vowel-consonant (VC), consonant-vowel-consonant (CVC), or other for all participants (n=7) and word pairs (n=1-4, variable by participant) for single syllable stimuli. **c.** The majority of electrodes (63%) show perceptual effects for only one stimulus, while 37% of electrodes encode the perceived word in 2 or more stimuli (1-syllable words only). Gray bars indicate multinomial distribution expected by chance (±s.d.).

